# Root-zone modelling frameworks identify this managed ecological interface as a predictor of pine seedling quality

**DOI:** 10.64898/2026.05.04.722574

**Authors:** Jamil Chowdhury, Nathan Milne, Melanie Wade, Bronwyn Thuaux, Phil Green, Ian Last, John Senior, Angus J. Carnegie, Ian C Anderson, Tarryn Turnbull, Krista L. Plett, Jonathan M Plett

## Abstract

Containerised forest nurseries increasingly supply planting stock for reforestation and plantation establishment, yet root-zone mycobiome assembly and functional importance during nursery production remain poorly integrated into assessments of seedling quality. We used operational radiata pine nursery production to test whether substrate amendment, crop-protection regime and fertigation intensity were associated with coordinated shifts in seedling performance and the root-associated mycobiome. Seedling height, root-collar diameter, root dry weight and mycorrhizal colonisation were integrated into a Composite Performance Index, and fungal communities were profiled using ITS2 sequencing, functional-guild annotation, variance partitioning and treatment-aware machine-learning models. Granular thiophanate-methyl/etridiazole was associated with the strongest improvement in composite seedling performance, whereas biochar shifted fungal community structure without improving growth, and reduced fertigation maintained seedling performance under the conditions tested. Across treatments, the root mycobiome retained a stable ectomycorrhizal-dominated backbone, but management and performance classes were associated with shifts among dominant ECM genera and lower-abundance saprotrophic, endophytic, mycoparasitic and pathogen-associated guilds. Mycobiome principal components explained additional variation in seedling performance beyond management treatments alone, and mycobiome-based classifiers separated high- and low-performing trays more effectively than management-only models. These findings position the containerised nursery plug as a managed ecological interface linking production inputs and root fungal community structure to seedling quality.

## 1. Introduction

Reforestation and establishment of plantation forestry depend on the reliable production of vigorous, disease-free planting stock, increasingly achieved in containerised nursery production that use intensive inputs such as controlled-release fertilisers, liquid fertigation, fungicide programs and substrate amendments (Carrillo et al., 2011; Rikala et al., 2004; Smaill et al., 2020; Veijalainen et al., 2007). Nursery performance is evaluated through visible traits, including shoot growth, root-collar diameter, and general plant health. However, these aboveground indicators may not fully capture the belowground biological condition of planting stock as nursery management also influences rooting and the root-zone mycobiome. Root-associated microbial communities play key roles in nutrient acquisition, stress tolerance, pathogen suppression and seedling establishment after outplanting into forest-establishment sites (Franco et al., 2014; Smaill and Walbert, 2013). Therefore, nursery management should be considered not only as a way to produce morphologically acceptable seedlings, but also as a process that conditions the microbial communities necessary to plant health.

Plant-soil feedback theory provides a useful framework for understanding these nursery-derived microbial legacies. This theory has been used to understand how plants modify the soil biotic community, and this modified community can then feedback positively or negatively on plant growth (Bever *et al*., 1997; van der Putten *et al*., 2013). Although plant–soil feedbacks are well established in natural and agricultural ecosystems (Ehrenfeld et al., 2005; Mariotte et al., 2018; van der Putten et al., 2013), they have rarely been treated as a manageable components or phenotype in containerised forest nurseries. This is important because the seedling plug is a confined root-zone micro-ecosystem. Within this confined environment, irrigation, fertiliser supply, fungicide exposure and substrate composition impose selection pressures on microbial assembly. The resulting plug-associated community is then carried with the seedling plug to the planting site where it could contribute to variation in disease tolerance and establishment outcomes of the tree host (Franco *et al*., 2014; Grove *et al*., 2019; Schneider *et al*., 2024).

The root fungal community of conifer seedlings contains several functional guilds with contrasting or complementary roles in these feedback processes. Ectomycorrhizal (ECM) fungi, often the most dominant microbial features of pine roots (El Karkouri *et al*., 2005; Richard *et al*., 2005; Smaill *et al*., 2020), form mutualistic symbioses that enhance nutrient and water uptake, improve stress tolerance, and can help protect against soil-borne pathogens by occupying infection sites or altering root exudation patterns (Colpaert et al., 1999; Parke et al., 1983; Sousa et al., 2012). Non-ECM root-associated fungi (for example, *Phialocephala, Meliniomyces, Trichoderma*, etc) are also common components of conifer roots and can provide similar benefits (Mandyam and Jumpponen, 2005; Rodriguez et al., 2009; Summerbell, 2005; Vohník et al., 2013; Wang et al., 2022). Saprotrophic fungi contribute indirectly by decomposing organic matter and recycling nutrients, and some genera can antagonise pathogens or influence plant defences (Asplund et al., 2019; Leifheit et al., 2024; Mašínová et al., 2017). Mycoparasitic fungi add another functional layer by attacking other fungi through mycoparasitism, competition or antibiosis (Geiger et al., 2022; Mascarin et al., 2022; Mukherjee et al., 2022; Singh et al., 2024; Yao et al., 2023). In contrast, pathogenic fungi such as *Fusarium, Phytophthora, Rhizoctonia* and *Dactylonectria* spp., directly damage roots and collars, causing damping-off, root rot or stem lesions that reduce seedling quality and survival (De Benedetti et al., 2025; Habibi et al., 2020; Lamichhane et al., 2017; Oxenham and Winks, 1963; Reglinski et al., 2009; Won et al., 2018). Thus, the balance among fungal guilds may be more informative than the abundance of any single taxon.

Previous studies show that conventional management can shift the microbiome balance in ecologically meaningful ways. In a *Pinus radiata* (radiata pine) bare-root nursery, Smaill and Walbert (2013) found that intensive fertilizer and fungicide use altered ECM community composition, promoting less-beneficial ECM taxa and reducing *Rhizopogon* species associated with better seedling nutrition. A subsequent multi-year field study demonstrated that radiata pine seedlings raised under reduced fungicide exposure had larger root collar diameters and higher survival for at least six years after planting compared with seedlings grown under standard fungicide regimes, suggesting that nursery chemical use can leave a detectable imprint on post-planting survival and forest-establishment outcomes (Smaill et al. 2020). Nutrient delivery can also act as a strong ecological filter. In containerised ECM-inoculated radiata pine, Stuart *et al*. (2023) showed that a mycocentric fertilisation regime increased belowground growth, mycorrhizal colonisation and root-tip nitrogen accumulation while shifting both ECM and non-ECM fungal community composition. Other biological or alternative inputs can also reshape nursery microbiomes and disease outcomes, including aerated compost tea against *Fusarium circinatum* in radiata pine (Otero et al. 2020) and amino-acid-based biostimulant fertigation that enriched root-associated fungi linked to growth-promoting functions (Chowdhury *et al*., 2026).

Despite the growing evidence, it remains unclear how root-zone nursery management practices interact to influence seedling quality and root mycobiome structure. Most nursery studies have examined individual management components, such as fertilisation, fungicide use, biological amendments or microbial inoculation, but operational nurseries apply these inputs simultaneously. This creates an important practical and ecological question whether seedling quality be improved by changing the root-zone environment rather than simply increasing nutrient input. We selected to test in a nursery environment three management axes – substrate amendment, crop-protection strategy and fertigation intensity - because each represents a practical way to modify the root-zone environment before seedlings are deployed for reforestation or plantation establishment. Substrate amendment with biochar was included because biochar can improve substrate structure, water retention and microbial habitat, although its effects are often context dependent (Jaafar et al., 2014; Kaudal et al., 2016; Knerr et al., 2025; Nemati et al., 2015). Granular thiophanate-methyl/etridiazole was incorporated into the substrate to test whether placing disease-suppressive active ingredients directly in the root zone can reduce early root disease pressure more effectively than relying only on later liquid crop-protection applications (Dumroese et al., 1990; Kataria and Grover, 1978; Krasnow and Hausbeck, 2017). Reduced fertigation was included to test whether moderate liquid nutrient input can maintain seedling quality while avoiding unnecessary input intensity and potentially supporting a more functionally balanced root mycobiome. From these, we used variance partitioning and machine-learning models to test whether root mycobiome composition added predictive information to seedling health and quality beyond management treatments and to identify fungal signatures associated with high- and low-quality planting stock. This study provides a transferable framework for microbiome-informed production of planting stock for reforestation, afforestation and plantation establishment.

## 2. Materials and methods

### 2.1 Experimental design, seedling growth conditions and nursery treatments applications

The experiment was conducted on container blocks at the Hancock Victorian Plantations (HVP) Gelliondale tree nursery in South Gippsland, Victoria, Australia. Seedlings of *radiata pine* were produced under commercial conditions using Transplant Systems 45-cell forestry trays, each cell holding approximately 93 cm³ of potting substrate. Seeds were soaked in water for 24 h, cold stratified at low temperature for 48 h, and then sown into a non-sterile potting medium based on composted pine bark.

For the substrate amendment treatments, three substrate mixes were made by combining the base composted pine bark with one of these amendment regimes: no amendment (SA-0), biochar addition (SA-BC), or a granular fungicide (SA-GF) that was a mix of Thiophanate-Methyl (50g/kg) and Etridiazole (30g/kg) incorporated into the substrate as a soil-applied fungicide (SA-GF). The biochar was a high-grade product produced from woody eucalypt feedstock at approximately 500 to 550°C. Biochar was incorporated into the medium prior to tray filling at 0.1 m³ per m³ of media. Granular fungicide was incorporated into the medium prior to tray filling at 300 g per m³ of media. After these amendments were blended thoroughly with the bark, each substrate mix received an identical basal fertiliser regime consisting of 2.5 kg m□³ of a 3–4 month slow-release fertiliser (NPK 18:2.6:9.9) and 5.5 kg m□³ of an 8–9 month controlled-release fertiliser (NPK 17:2.3:10; Agsolutions Pty Ltd, Sydney, Australia). This ensured that all substrate treatments received the same controlled-release fertiliser loading, and that the only difference among mixes at sowing was the presence or absence of biochar or granular fungicide. Seeds were then sown directly into these amended and fertilised mixes. A starter fertigation (Peters Professional, 1 g L□¹) was applied uniformly to all seedlings for the first six weeks after germination to promote even establishment.

The tray-based trial was designed to match with operational nursery practice. Trays were grouped into plots of 36 trays, with each plot placed on a separate bench and allocated a single crop-protection regime to avoid cross-contamination among fungicide programs. Four plots (each representing a distinct crop-protection regime) constituted one replication, and this structure was repeated across six replications for a total of 38,880 seedlings. Within each replication, the four crop-protection treatments were randomly assigned to benches and comprised: no fungicide (CP-Nil), the nursery standard fungicide program (CP-Std), a liquid fungicide -based chemical fungicide program (CP-FunA), and a *Trichoderma*-based biological control program (CP-Trich). In CP-FunA, liquid fungicide containing a mixture of Thiophanate-Methyl (250g/kg) and Etridiazole (150g/kg) was applied fortnightly according to the nursery program. In CP-Trich, a commercially available product containing *Trichoderma harzianum* was applied during week 1 after sowing and then monthly, with the *Trichoderma* formulation applied at 150 g per 15 L water using a knapsack. In CP-Std, the standard nursery fungicide regime was applied weekly and reduced as required according to routine nursery practice. All crop-protection applications were implemented by nursery staff using their routine spray or drench schedules for each program to reflect operational conditions.

Within each crop-protection plot, trays were allocated to the three substrate mixes (SA-0, SA-BC, SA-GF), and tray positions were randomised within plots to minimise within-bench environmental bias. Three randomly assigned plots were further subjected either 100% of the recommended liquid fertiliser concentration (FR-100) or 75% of the recommended concentration (FR-75). Weekly fertigation was delivered using a Seltec liquid fertiliser (50 mL per 15 L water per 1000 seedlings) through the overhead boom irrigation system, while day-to-day irrigation followed the nursery’s standard automated watering schedule. Border trays were installed as buffer rows along exposed bench edges and were excluded from all measurements. Seedlings were grown under ambient nursery conditions (HVP Plantations, Victoria, Australia) and seedling growth, health, and root traits were assessed 12 months after sowing, coinciding with the operational pre-dispatch stage for planting stock.

### 2.2 Morphological and physiological assessments of samples

Seedling morphological parameters including plant height, root-collar diameter, and root dry weight were assessed on tall rays at 12 months after sowing. Visual estimation of mycorrhizal colonisation as the mean proportion of colonised root tips and plug integrity were also recorded on a subset of plants for each blocked replicate. To obtain a single quantitative measure of overall seedling performance, we calculated the Composite Performance Index (CPI) as an unweighted composite index by standardising each tray-level trait to a z-score and averaging z-scores across traits, an approach commonly used to combine multiple performance variables measured on different scales into a single index (Hooper and Vitousek, 1998; Hu et al., 2025; Maestre et al., 2012; Nunes et al., 2025). For each trait *T* in the set (plant height, root-collar diameter, root dry weight, and mycorrhizal score) a z-score was calculated for tray *i* as

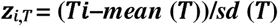

where T*_i_* is the mean value of trait *T* for tray *i*, *mean (*T) is the overall mean of that trait across all trays, and *sd* (T) is its standard deviation. The CPI for tray *i* was then defined as the arithmetic mean of the four-trait z-scores. Therefore, positive values indicate above-average performance across height, stem thickness, root biomass, and mycorrhizal colonisation while negative values indicate below-average performance. This CPI was used as the response variable in subsequent variance partitioning and machine-learning analyses.

### 2.3 DNA extraction, amplicon sequencing and taxonomic assignment

Approximately 250 mg of lateral roots taken randomly from the root balls of three plants per replication were collected, combined, and frozen for DNA extraction. In total, six separate block replicates per treatment were extracted and prepared for community sequencing (n=6). DNA was extracted using the DNeasy PowerSoil Pro Kits (Qiagen) following the manufacturer’s protocol with the addition of a cryo-grinding step at the beginning to ensure efficient disruption and homogenisation of the root tissues. Following Chen et al. (2021), the ITS region of fungi was amplified using a two-step PCR protocol for Illumina MiSeq Meta-barcoding, with 20 ng of DNA and the primers Forward: 5’-CCTACACGACGCTCTTCCGATCTNNNNGTGARTCATCGAATCTTTG-3’ and Reverse: 5’-GTGTGCTCTTCCGATCTCCTCCGCTTATTGATATGC-3’. Samples were multiplexed and sequenced on the Illumina MiSeq platform at the Advanced Gene Technology Centre, New South Wales Department of Primary Industries and Regional Development. DNA extraction blanks controls were processed in parallel with samples. All blank controls yielded < 200 raw reads and < 10 post-filter reads, and shared no unique ASVs with root samples; consequently, they were excluded from the final phyloseq object.

### 2.4 Mycobiome community analyses

Amplicon data processing and community analyses were conducted in R (R Core Team; https://www.r-project.org/) using the packages dada2 (Callahan *et al*., 2016), ShortRead, Biostrings, phyloseq (McMurdie and Holmes, 2013), vegan (Oksanen *et al*., 2019), and tidyverse (Wickham *et al*., 2019). Illumina ITS2 reads were first trimmed of primer sequences using Cutadapt, with forward and reverse primers provided in both orientations so that primer motifs were removed wherever they occurred in the reads. Quality filtering and denoising followed the standard dada2 pipeline (Callahan *et al*., 2016): low-quality reads and reads containing ambiguous bases were discarded, error models were learned from the data, and amplicon sequence variants (ASVs) were inferred and de-replicated. Chimeric sequences were removed using the consensus method implemented in removeBimeraDenovo, and the resulting non-chimeric sequence table was retained for all downstream analyses.

Taxonomy was assigned to each ASV using assignTaxonomy against the UNITE fungal reference database (dynamic release v9, release date 2023-07-18, DOI: 10.15156/BIO/2938066). A phyloseq object was then created by combining the ASV table, the taxonomy table, the sample metadata and the DNA sequences for each ASV stored in the refseq slot. ASVs were renamed with short “ASV” identifiers for readability while maintaining a key to the underlying sequences. Treatment effects of substrate amendment on alpha diversity were evaluated using analysis of variance with substrate amendment as a fixed factor, followed by Tukey’s HSD tests where appropriate.

To relate community composition to taxonomic and functional structure, ASV counts were agglomerated to genus level using tax_glom, and genus-level relative abundances were computed per sample. For graphical summaries, relative abundances were averaged within treatment combinations and low-abundance genera were grouped into an “Other” category to improve legibility. Genera were then annotated using the FungalTraits database (Põlme *et al*., 2021) and assigned to broad lifestyle categories such as ectomycorrhizal symbionts, root endophytes, saprotrophs, potential plant pathogens and so on. Guild-level relative abundances were obtained by summing genera within each lifestyle class and visualised as stacked bar plots to highlight how substrate amendment and associated nursery treatments redistributed functional guilds in the root mycobiome. For later modelling steps, the amplicon table was further collapsed to genus level and transformed to centred log-ratio (CLR) coordinates. ASV counts were summed within each genus for each sample and counts were log-transformed and centred by subtracting the sample-wise mean log abundance. The resulting CLR-transformed genus abundances were appended to the sample metadata and used as continuous predictors in variance partitioning and machine-learning analyses.

### 2.5 Quantifying management and mycobiome contributions to CPI using bootstrap linear models

To measure the impact of root mycobiome composition and nursery management levers on seedling performance, we used a linear model framework with bootstrap uncertainty to break down explained variation in the CPI. The taxonomy table was used to collapse the root mycobiome data from ASVs to genus, which was then handled as compositional data. In accordance with standard procedure for log-ratio analysis of compositional sequencing data, genus counts were transformed using a centered log-ratio (CLR) after adding a pseudocount of 1 to handle zeros (Aitchison, 1982; Gloor et al., 2017, 2016). Principal components analysis (PCA) was then applied to the CLR genus matrix to obtain a compact description of community composition where the first three PCs were retained as the microbiome predictor set (Jolliffe, 2002). Following this we constructed and compared a management-only model (CPI ∼ Substrate amendment + Crop protection regime + Fertigation) to a management + microbiome model that added CLR_PC1–CLR_PC3. Model fit was summarised using R², adjusted R², and RMSE. For every predictor block (Microbiome, Substrate amendment, Crop protection regime, Fertigation), we calculated two complementary quantities in order to attribute explained variation at the block level. First, the block-level unique contribution (semi-partial R²; ΔR²) was calculated as the reduction in full-model R² when the block was removed while keeping all other predictors. Second, the block-only association was calculated as the R² from a model containing only that block. Nonparametric bootstrap resampling (500 iterations) was used to quantify uncertainty (Efron and Tibshirani, 1994).

### 2.6 Machine-learning detection of performance-associated mycobiome signatures

To test whether root mycobiome composition carried a diagnostic signal of planting-stock quality, we trained supervised machine-learning classifiers to separate high- and low-CPI trays. CPI values were centred and scaled to obtain CPI_z. Extreme performance classes were then defined using the upper and lower tertiles of CPI_z. Trays in the upper tertile were labelled as Top-CPI, trays in the lower tertile were labelled as Bottom-CPI, and middle-tertile trays were excluded to maximise contrast between classes. We compared three predictor sets: management-only, mycobiome-only and combined management plus mycobiome predictors. The management-only set included substrate amendment, crop-protection regime and fertigation rate, encoded as treatment-level dummy variables. The mycobiome-only set included genus-level centred log-ratio (CLR) transformed fungal abundances. The combined set included both treatment-level management predictors and genus-level CLR fungal abundances. This structure allowed us to test whether fungal profiles classified Top- and Bottom-CPI trays better than treatment labels alone. Random Forest classifiers were fitted using the *ranger* package (Breiman, 2001; Wright and Ziegler, 2017) and gradient-boosted tree classifiers were fitted using *xgboost* (Chen and Guestrin, 2016). Model performance was assessed using repeated stratified five-fold cross-validation, preserving the Top-CPI/Bottom-CPI ratio within each fold. For each resample, models were trained on four folds and tested on the held-out fold. Out-of-fold predictions were used to calculate receiver operating characteristic curves, area under the curve (AUC), accuracy, sensitivity, specificity and balanced accuracy (Robin et al., 2011). Feature importance was calculated separately for each predictor set and algorithm. For Random Forest, permutation importance was used; for XGBoost, gain-based importance was used (Chen and Guestrin, 2016). Because these importance measures are on different scales, values were normalised within each algorithm and predictor set to a 0–100 scale before comparison. To examine the direction of the strongest fungal associations, partial dependence profiles were generated for leading fungal predictors from the combined model using the *pdp* framework (Greenwell, 2017). These profiles described how the predicted probability of Top-CPI classification changed across the observed CLR abundance range of each genus, while averaging over the remaining predictors.

## 3. Results

### 3.1 Root-zone substrate amendment had the strongest effect for improving multi-trait seedling quality

Our data reveal that across all seedling traits, substrate amendment (SA) showed the strongest association with planting-stock performance, with crop protection (CP) and fertigation regime (FR) acting as secondary modifiers. Relative to the unamended control substrate (SA-0), root-zone incorporation of the granular thiophanate-methyl/etridiazole amendment (SA-GF) was associated with higher values for the three core nursery quality traits which are plant height, root-collar diameter and root dry weight (Figure 1, Table S1, S2). On average, SA-GF significantly increased seedling height by around 10 mm and root-collar diameter by 4 mm, while root dry weight increased by roughly 25% compared with the unamended control substrate. The Biochar-amended substrate (SA-BC) produced relatively smaller and statistically insignificant increases in those traits. Plug integrity scores and visual mycorrhizal colonisation did not differ meaningfully among substrate treatments at the main-effect level.

**Figure 1:**
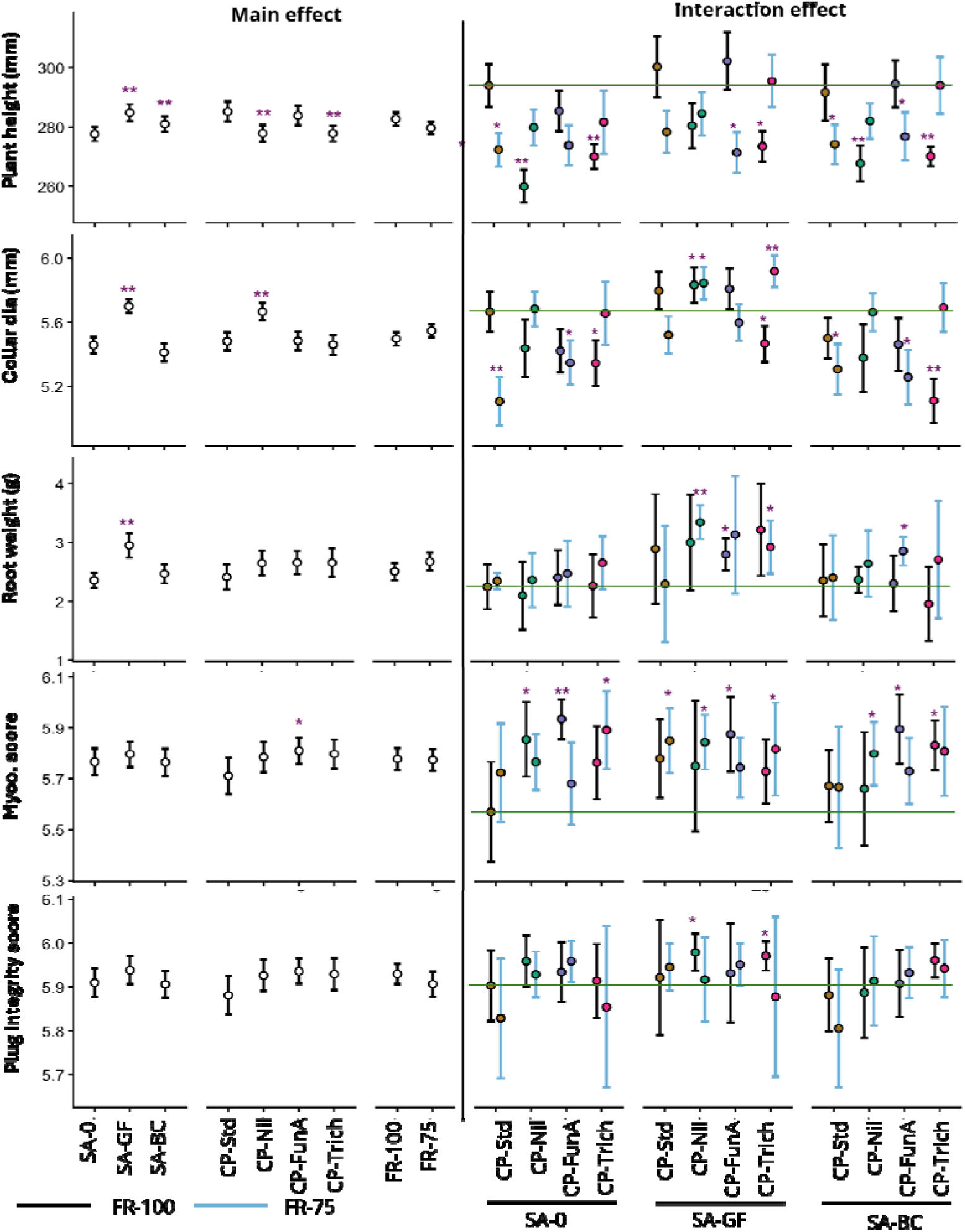
Main and interaction effects of substrate amendment, crop protection regime and fertigation regime on radiata pine seedling morphology at nursery dispatch before outplanting. (a) The left-hand panels show main effects of substrate amendment (SA-0, SA-GF, SA-BC), crop protection regime (CP-Std, CP-Nil, CP-FunA, CP-Trich) and fertigation rate (FR-100, FR-75) on key pine seedling growth parameters. (b) The right-hand panel shows the full set of interaction combinations, faceted by substrate, with crop protection treatments on the x-axis. Points represent treatment means and vertical bars show 95 percent confidence intervals based on row-level data. Black symbols and error bars denote FR-100, and blue symbols denote FR-75; point fill colour indicates crop protection regime. The horizontal green line marks the mean of the benchmark nursery combination (SA-0 / CP-Std / FR-100). Asterisks indicate significant differences from the relevant control (SA-0 for substrate, CP-Std for crop protection, FR-100 for fertigation, and SA-0 / CP-Std / FR-100 for interaction combinations) after Holm adjustment for multiple comparisons (* p < 0.05, ** p < 0.01).

Crop protection regime also affected plant height and root-collar diameter, although their effects were smaller than those of substrate amendment. Removing fungicide inputs (CP-Nil) or replacing the standard regime with a *Trichoderma*-based program (CP-Trich) reduced plant height by approximately 2-3 percent compared to the standard fungicide program (CP-Std) as the reference. In contrast, CP-Nil treated seedlings had approximately 3% thicker collar diameter than those under CP-Std. Seedlings treated with the liquid thiophanate-methyl/etridiazole program (CP-FunA) maintained plant growth close to the standard program. Root dry weight, plug integrity and mycorrhizal scores were only weakly affected by crop protection regime at the main-effect level. Reducing fertigation from the standard rate (FR-100) to 75% of the recommended rate (FR-75) did not produce a significant negative effect on seedling growth traits at the main-effect level, indicating that reduced liquid nutrient input maintained planting-stock performance in this experiment (Figure 1).

The three-way interaction plots and CPI ranking further showed how the management components performed in combination (Figure 2). The benchmark nursery combination, comprising unamended substrate, standard crop protection and full fertigation (SA-0 / CP-Std / FR-100) was positioned near the centre of the CPI distribution. This indicates that the benchmark program produced seedlings of approximately average composite performance when height, root-collar diameter, root dry weight and mycorrhizal score were integrated into a single index. In contrast, many of the highest-ranked combinations shared the granular thiophanate-methyl/etridiazole substrate amendment (SA-GF), often in combination with either no fungicide or Trichoderma-based crop protection and reduced fertigation. At the lower end of the CPI spectrum, the poorest-performing treatments tended to involve either the unamended control substrate or the biochar-amended substrate, with crop-protection regime or fertigation rate playing a smaller role. Although only a subset of specific trait combinations showed statistically significant deviations from the control in the interaction panels, the overall pattern is consistent with root-zone delivery of the granular thiophanate-methyl/etridiazole amendment being the strongest tested intervention associated with higher multi-trait seedling quality. Fertigation rate and crop-protection regime appeared to modify this response, either reinforcing or reducing the gains depending on the treatment combination (Figure 2).

**Figure 2:**
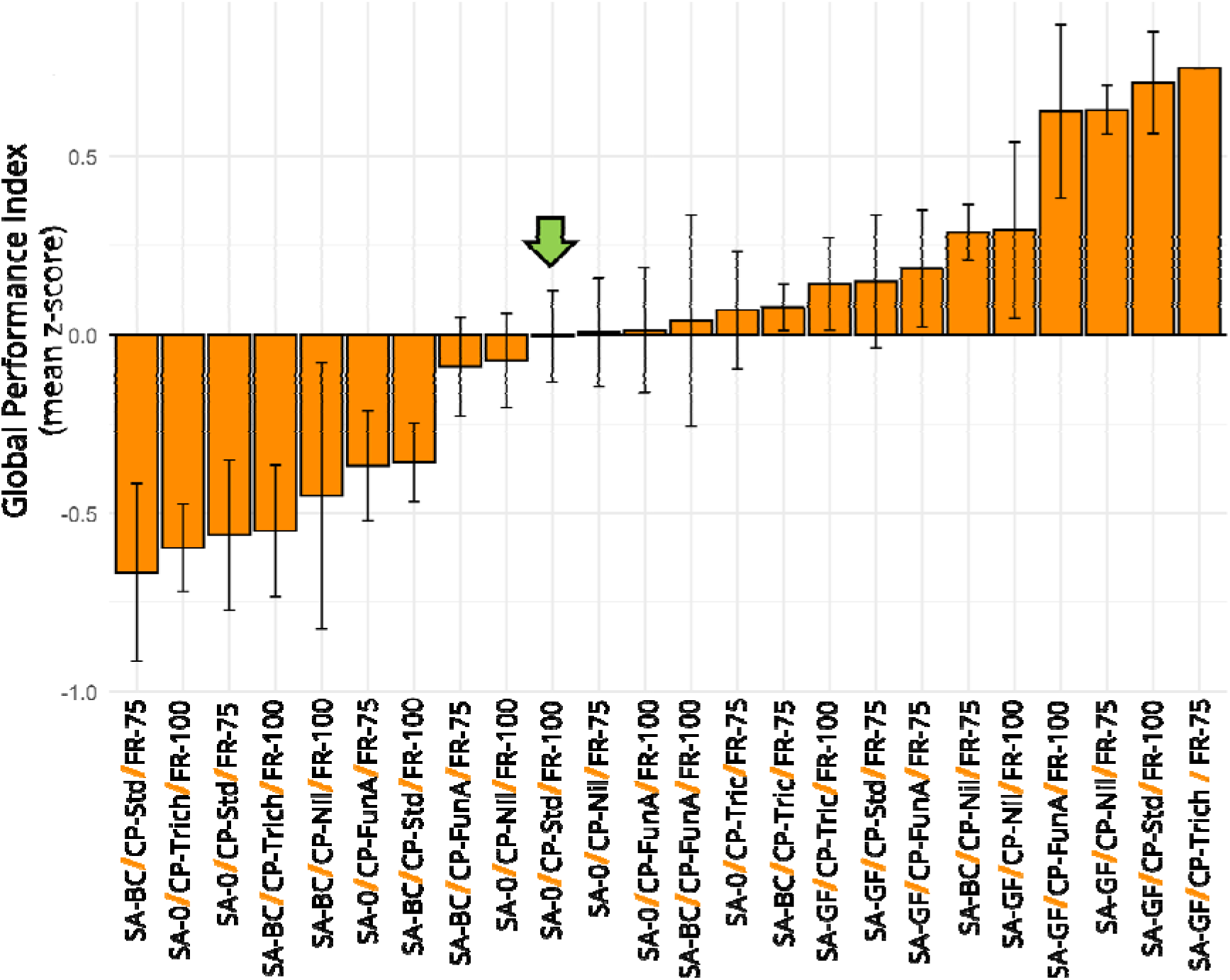
Substrate amendment with granular fungicide (SA-GF) was associated with higher composite seedling performance. Composite performance index (CPI) ranking of all substrate amendment × crop protection × fertigation combinations. Bars show the mean CPI for each treatment combination, expressed as the mean z-score across four nursery quality traits (height, collar diameter, root dry weight and mycorrhizal score). Values above zero indicate better-than-average composite performance and values below zero indicate poorer-than-average performance. The green arrow marks the benchmark nursery combination (SA-0 / CP-Std / FR-100), which lies near the centre of the distribution. Error bars indicate standard error.

### 3.2 Management regimes rebalanced functionally informative root fungal guilds beneath a stable ectomycorrhizal backbone

Across all treatments, the root-associated fungal community retained a strong ectomycorrhizal (ECM) backbone, dominated by *Rhizopogon*, *Thelephora*, *Wilcoxina*, *Suillus*, *Inocybe* and *Serendipita*. Combined, these genera accounted for about 91–94% of all reads across the experiment (Figure 3, S1, S2). Against this stable ECM background, substrate amendment (SA-0, SA-GF, SA-BC) rebalanced the lower-abundance, but functionally informative guilds, as assigned to genera using the FungalTraits database (Põlme et al., 2021); Figure 3). Relative to the unamended control (SA-0), root-zone incorporation of the granular thiophanate-methyl/etridiazole amendment (SA-GF) reduced the ECM guild only slightly (−3.1%), but substantially shifted the relative abundance of individual ECM genera. For example, it reduced the relative abundance of *Rhizopogon* and *Inocybe* (-12%, -36%, respectively vs SA-0) and increased the relative abundance of *Serendipita* and *Pustularia* (+65%, +190%, respectively vs SA-0; Figure 3).

**Figure 3.**
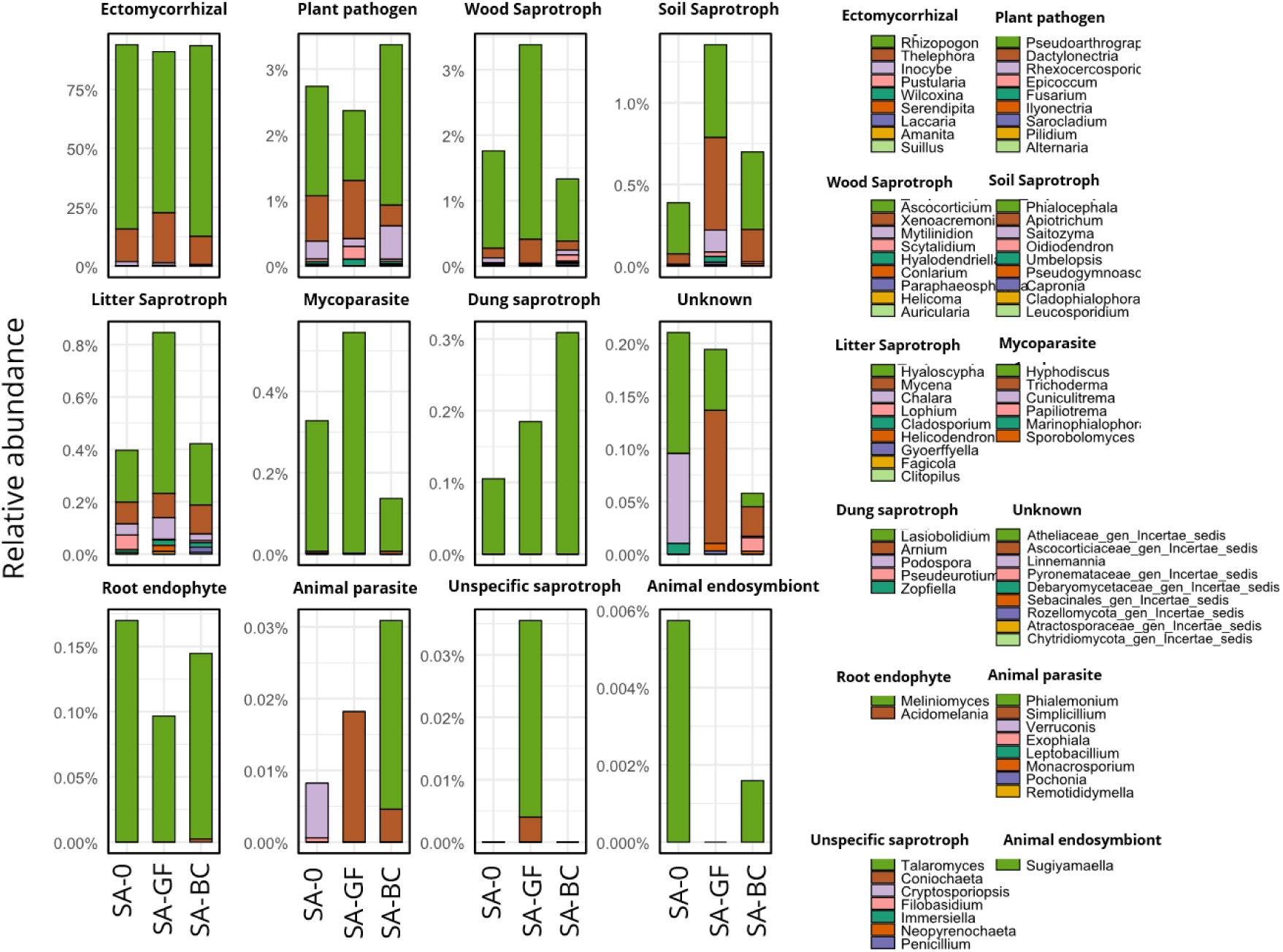
Functional guild structure of the root-associated mycobiome across substrate amendment treatments. Relative abundance of fungal functional guilds and dominant genera in 12-month radiata pine seedlings grown under three substrate amendments: control substrate with no amendment (SA-0), Granular fungicide amendment (SA-GF) and biochar amendment (SA-BC). Bars show the proportion of reads per treatment that belong to each guild, faceted by guild. Within each panel, colours represent the top-ranked genera plus an “Others” category. Functional guilds were assigned to genera using the FungalTraits database.

SA-GF also strongly enriched saprotrophic and mycoparasites, a guild antagonistic to pathogens. Wood saprotrophs (e.g. *Ascocorticium, Xenoacremonium*) increased by ∼92%, soil saprotrophs (e.g. *Apiotrichum, Phialocephala*) by ∼2.5-fold, litter saprotrophs (e.g. *Mytilinidion, Hyalodendriella*) by 113.6% and mycoparasites (e.g. *Hyphodiscus, Trichoderma*) by ∼66% compared with SA-0, while root endophytes (e.g. *Meliniomyces, Acidomelania*) declined (−43.1%). The potential plant-pathogen guild (e.g. *Pseudoarthrographis, Dactylonectria, Fusarium*) showed a modest decrease in SA-GF (−13.6%) and an increase in SA-BC (+23.1%). Among soil saprotrophs, *Apiotrichum* showed the clearest response, increasing nearly eight-fold in SA-GF (+793.0%) and approximately two-fold in SA-BC (+211.0%), while *Phialocephala* increased more moderately (+81.4% and +51.7%, respectively; Figure 3). Changes in the CP regime (CP-Std, CP–Nil, CP–FunA, CP–*Trichoderma*) produced relatively smaller shifts than substrate amendment, indicating that crop protection regime acted mainly as a secondary filter on fungal assembly (Figure S1). The overall ECM fraction and its dominant genus (*Rhizopogon*) remain nearly unchanged. However, relative abundance of ECM genera *Inocybe, Thelephora* and *Wilcoxina* were considerably increased when CP-Std was replaced by any of CP–Nil, CP–FunA, CP–*Trichoderma* (Figure S1). Crop-protection effects were therefore concentrated in lower-abundance guilds rather than in the ECM backbone. Relative to the standard programme (CP-Std), CP–Nil reduced potential plant-pathogen guild (−18.9%) and wood and litter saprotrophs (−22.5% and −31.8%) but strongly increased root endophytes (+70.6%), whereas CP–FunA and CP–Trichoderma generally reduced saprotrophs and mycoparasites (e.g. wood saprotrophs −55.0% and −41.9%; mycoparasites −35.9% and −55.6%) and also lowered root endophytes (−60.1% and −67.4%), while keeping potential plant-pathogen guild close to the standard level (+9.9% and +5.7%). Within soil saprotrophs (e.g. Apiotrichum, Phialocephala), Apiotrichum again showed a strong directional response, being enriched under CP–Nil and CP–Trichoderma (+216.3% and +203.9% vs standard).

Adjusting liquid fertigation from 100% (FR-100) to 75% (FR-75) of the recommended rate had only modest effects on overall community structure with the ECM guild and its dominant genera essentially unchanged (Figure S2). However, reduced fertigation rebalanced several lower-abundance guilds. Under FR-75, the potential plant-pathogen guild (e.g. *Pseudoarthrographis, Dactylonectria*) increased by +31.6%, soil and litter saprotrophs (e.g. *Apiotrichum, Phialocephala, Ascocorticium*) by +23.5% and +9.9%, and mycoparasites (e.g. *Hyphodiscus, Trichoderma*) by +85.6%. Root endophytes (e.g. *Meliniomyces*) declined sharply (−63.5%). Within the soil saprotrophs, *Phialocephala*, a common dark septate root endophyte of conifers, increased by 120% under reduced fertigation, consistent with a shift in decomposer–endophyte balance under reduced liquid nutrient input.

### 3.3 Root mycobiome composition and substrate amendment are the dominant drivers of variation in seedling performance

To quantify the relative contribution of nursery management blocks and root mycobiome composition to the performance of seedlings, we partitioned the explained variations in the CPI using a bootstrap linear model variance framework. Adding root mycobiome composition (CLR_PC1-PC3) to the management-only model substantially improved model performance, increasing the adjusted R² from 0.225 to 0.447 and lowering the RMSE from 0.547 to 0.437 (Figure 4a, Table S3). Thus, inclusion of the mycobiome block nearly doubled the proportion of CPI variation explained by the model. Bootstrap resampling indicated moderate uncertainty around the absolute effect size, but the full model consistently retained greater explanatory power across resamples, as reflected by the bootstrap confidence interval for R² (Table S4).

**Figure 4.**
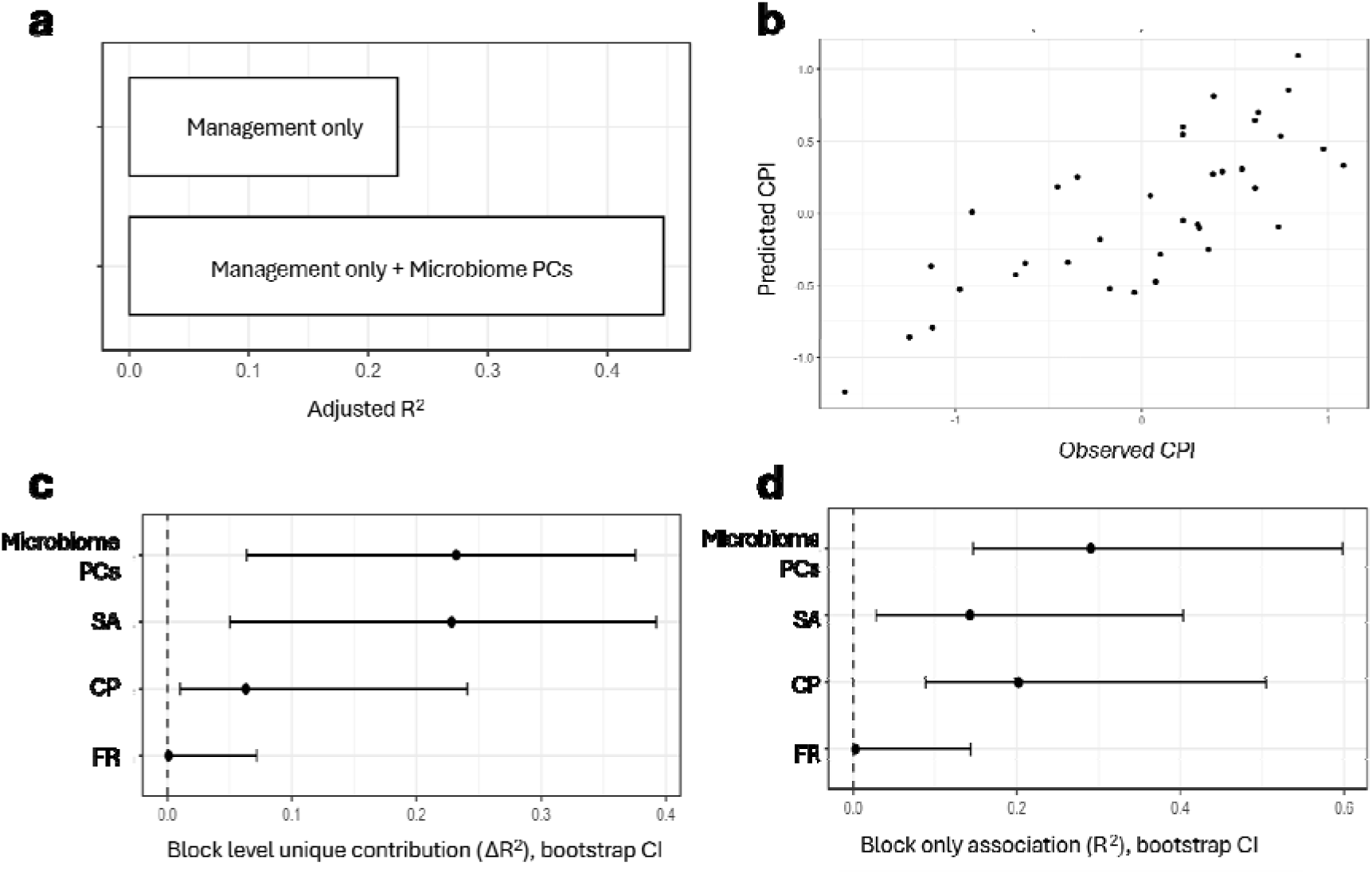
Linear model variance decomposition of CPI using nursery management blocks and mycobiome PCs. (a) Adjusted R² comparison between a management-only model and the model including root mycobiome PCs (CLR_PC1–PC3). (b) Predicted versus observed CPI for the full model. (c) Block-level unique contribution to explained variance (ΔR²), defined as the reduction in model R² when each block is removed while all other blocks are retained. (d) Block-only association with CPI (block-only R²), calculated from models containing each block alone. Error bars show 95% bootstrap confidence intervals (CI). Points show point estimates and horizontal lines show 95% bootstrap confidence intervals (500 resamples).

We then examined the contributions at the block level in two complementary ways. First, we estimated the unique contribution of each block as the additional variance explained after accounting for overlap with the remaining predictors (incremental gain). Under this criterion, root mycobiome composition and substrate amendment made the largest non-overlapping contributions to explained variation in CPI, whereas crop-protection regime added a smaller additional component and fertigation contributed little unique variance within this dataset (Figure 4c; Table S5). Second, we examined block-only associations, in which each block was assessed separately to quantify how much variance it explained on its own without any other predictor. In these single block models, the mycobiome block showed the strongest standalone association with CPI, while crop-protection regime also showed a moderate association, consistent with shared explanatory structure between management treatments and the mycobiome (Figure 4d; Table S6). These analyses indicate that root mycobiome profiles contained structured information about planting-stock quality beyond treatment categories alone (Romero et al., 2025).

### 3.4 Machine-learning benchmark identified predictive fungal signatures of extreme composite seedling performance

Given that root mycobiome composition explained additional variation in CPI, we next tested whether high- and low-performing trays differed in fungal community profiles associated with performance. Trays in the upper and lower CPI tertiles were classified as Top-CPI and Bottom-CPI groups, respectively, and the middle tertile was excluded to maximise contrast between performance classes. We then trained binary machine-learning classifiers using three predictor sets: management-only predictors, mycobiome-only predictors and combined management plus mycobiome predictors. Management variables were one-hot encoded at the treatment level so that Random Forest and XGBoost evaluated the same treatment-level predictor structure (Fig. 5a,b; Table S7; Table S8).

**Figure 5.**
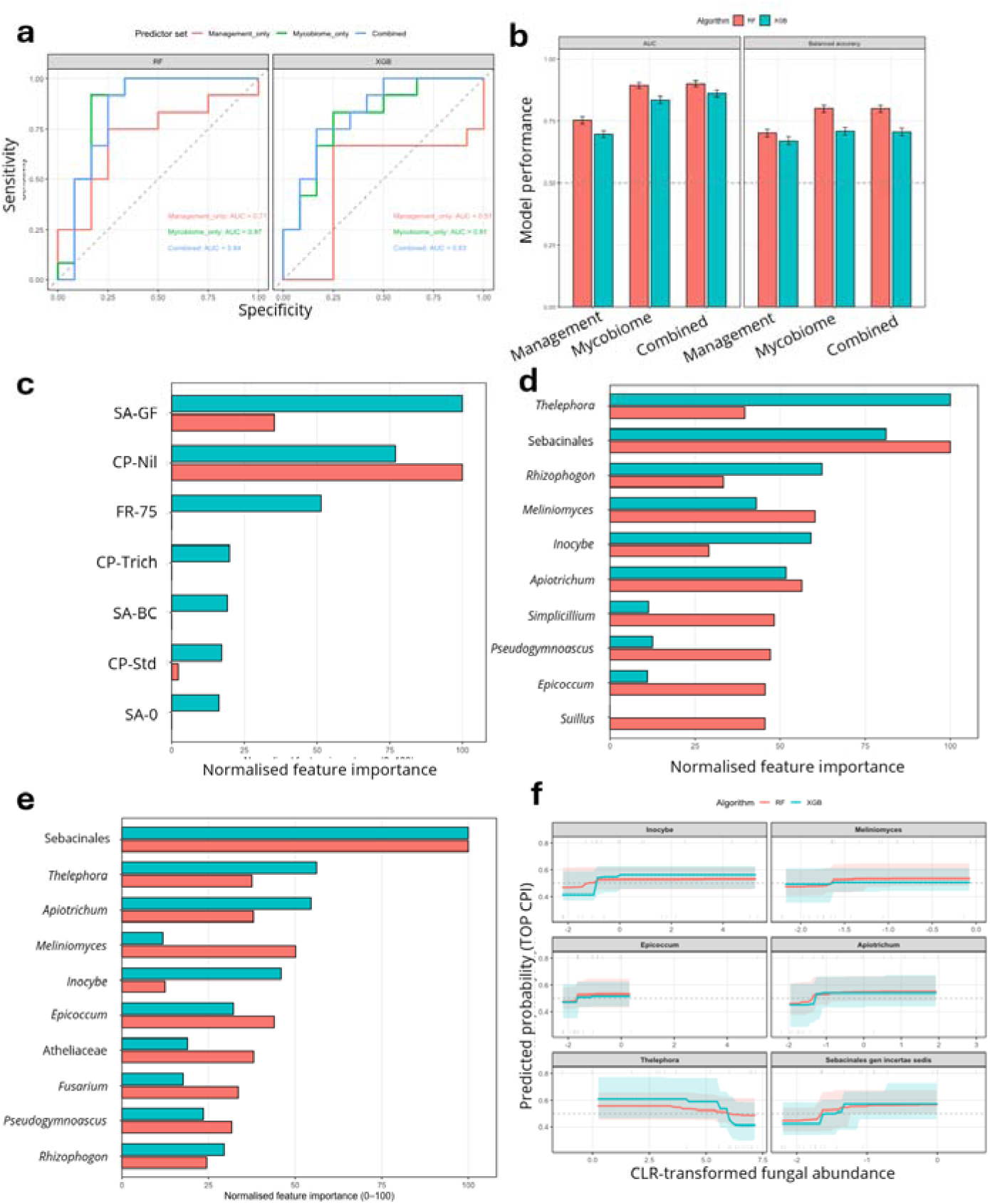
Treatment-aware machine-learning benchmark for classification of high- and low-performing nursery trays. Trays in the upper and lower tertiles of the Composite Performance Index (CPI) were classified as Top-CPI and Bottom-CPI groups, respectively, and the middle tertile was excluded to maximise contrast between performance classes. Models were trained using three predictor sets: management-only predictors, mycobiome-only predictors and combined management plus mycobiome predictors. Management predictors were one-hot encoded at the treatment level, and mycobiome predictors were genus-level centred log-ratio (CLR) transformed fungal abundances. (a) Out-of-fold receiver operating characteristic (ROC) curves for Random Forest (RF) and XGBoost (XGB) classifiers. Curves show predicted probabilities for the Top-CPI class, with AUC values shown within each panel. (b) Cross-validated model performance across repeated stratified folds. Bars show mean ± SE for AUC and balanced accuracy. (c) Top treatment-level predictors from the management-only models. (d) Top fungal predictors from the mycobiome-only models. (e) Top predictors from the combined management plus mycobiome models. (f) Partial dependence plots for leading fungal predictors from the combined model. Lines show the model-predicted probability of Top-CPI classification across the observed CLR abundance range for each genus, shaded bands show percentile intervals from bootstrap model refits, and rug marks show observed sample values. Feature-importance values in panels c–e were normalised within each algorithm to a 0–100 scale and should be interpreted as relative ranks within each model. Partial dependence curves indicate model-based directional associations and should not be interpreted as causal effects.

Management-only classifiers showed weaker discrimination between Top- and Bottom-CPI trays than classifiers containing mycobiome predictors. In the out-of-fold ROC analysis, the management-only model gave an AUC of 0.71 with Random Forest and 0.51 with XGBoost (Fig. 5a; Table S7). By contrast, mycobiome-only models gave higher AUC values of 0.87 and 0.81, respectively, while combined models gave AUC values of 0.84 and 0.83 (Fig. 5a; Table S7). The repeated cross-validation summary showed the same overall pattern, with higher mean AUC and balanced accuracy for models containing mycobiome predictors than for management-only models (Fig. 5b; Table S8). This indicates that fungal community composition provided stronger diagnostic information about CPI class than treatment labels alone.

Treatment-level feature importance confirmed that management predictors still carried biologically meaningful information. In the management-only models, the granular thiophanate-methyl/etridiazole substrate amendment, the no-fungicide crop-protection regime and reduced fertigation were among the most informative treatment-level predictors, although their relative ranking differed between Random Forest and XGBoost (Fig. 5c; Table S9). This shows that substrate amendment, crop-protection regime and fertigation rate contributed to performance-class separation. However, when management and mycobiome predictors were included together, the top-ranked predictors were dominated by fungal features rather than management variables (Fig. 5e; Table S9).

The mycobiome-only models repeatedly selected ECM and root-associated fungal genera as informative predictors (Fig. 5d; Table S9). These included ECM genera such as *Thelephora, Rhizopogon, Inocybe* and *Suillus*, which are consistent with the central role of ectomycorrhizal fungi in conifer nutrition, root function and stress tolerance (Colpaert et al., 1999; Parke et al., 1983). Sebacinales gen. incertae sedis and *Meliniomyces* also appeared among the most informative root-associated fungal predictors. In addition, the models selected taxa with saprotrophic, endophytic or antagonistic associations, including *Apiotrichum, Epicoccum, Simplicillium* and *Pseudogymnoascus*, consistent with the broader role of non-ECM fungi in decomposition, root-zone nutrient cycling and fungal interactions (Asplund et al., 2019; Leifheit et al., 2024; Mašínová et al., 2017). In the combined model, the most informative predictors again remained mainly fungal, with Sebacinales gen. incertae sedis, *Thelephora, Apiotrichum, Meliniomyces, Inocybe, Epicoccum*, Atheliaceae gen. incertae sedis, *Fusarium, Pseudogymnoascus* and *Rhizopogon* among the top features (Fig. 5e; Table S9, sheet Importance_all). The presence of *Fusarium* among the selected predictors indicates that both mutualistic or potentially beneficial fungal lineages and pathogen-associated lineages contributed to the performance-associated mycobiome signal, consistent with guild assignments based on the FungalTraits database (Põlme et al., 2021).

To examine the direction of the strongest fungal associations in the final model, we used partial dependence plots for leading fungal predictors from the combined classifier (Fig. 5f). Higher CLR abundance of Sebacinales gen. incertae sedis, *Apiotrichum, Inocybe, Epicoccum* and *Meliniomyces* generally increased the predicted probability of Top-CPI classification within the observed abundance range. *Thelephora* showed a more complex modelled response, with predicted Top-CPI probability declining at the upper end of its CLR range (Fig. 5f). These partial-dependence curves provide model-based directional associations, but they should not be interpreted as causal effects because fungal predictors are compositional and correlated. Together, the machine-learning benchmark shows that root mycobiome profiles carried diagnostic information associated with planting-stock quality beyond treatment labels alone.

### 3.5 Guild-level rebalancing within an ECM-dominated root mycobiome differentiated high-and low-performing nursery trays

To place these machine-learning signals into a broader guild-level context, we compared the relative abundance of all detected genera between Bottom and Top CPI tiers after assigning guilds using the FungalTraits database (Põlme *et al*., 2021). Across the whole community, ECM fungi remained overwhelmingly dominant in both tiers (c. 93% of reads), indicating that high- and low-performing trays shared the same ECM-dominated backbone (Fig. 6a, Table S10). However, the composition of this ECM backbone differed strongly between performance classes. Rhizopogon remained the dominant ECM genus in both tiers, but its relative abundance declined from 86.73% of all reads in Bottom-CPI trays to 70.83% in Top-CPI trays. In contrast, *Thelephora* increased from 6.54% in Bottom-CPI trays to 20.83% in Top-CPI trays. Inocybe was also detected only in Top-CPI trays, where it accounted for 1.23% of reads. High-performing trays contained a larger share of wood and soil saprotrophs (from 2.5 to 3.1% and 0.5 to 0.8% of reads, respectively), mycoparasites (0.2 to 0.4%) and root endophytes (0 to 0.1%), while low-performing trays carried slightly higher fractions of plant pathogens, dung saprotrophs and unassigned taxa. These patterns show differences in the relative contribution of decomposer, antagonistic and endophytic guilds within an otherwise ECM-dominated community.

**Figure 6:**
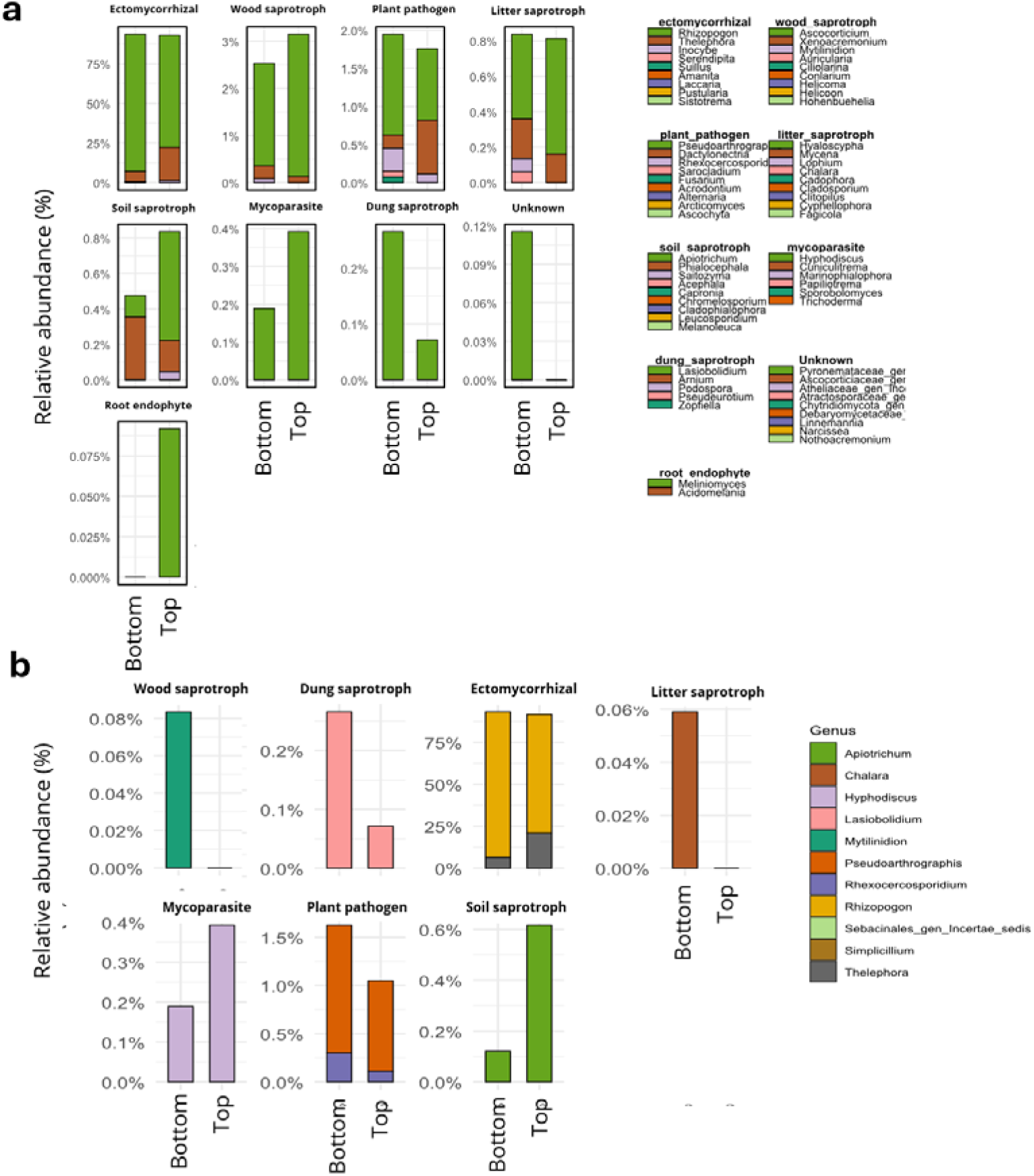
Top- and bottom-performing trays shared an ECM-dominated backbone but differed in the relative contribution of saprotrophic, antagonistic and pathogen-associated guilds. Relative abundance of fungal guilds in Bottom- and Top-CPI trays after genus-level assignment with the FungalTraits database. **(a)** Relative abundance of fungal functional guilds and dominant genera in Bottom and Top performance tiers (PerfTier), based on FungalTraits guild assignments. Each bar represents the average relative abundance within a tier, separated by guild. Within each facet, stacked colours represent the main contributing genera plus an “Others” category. **(b)** Relative abundance of ML-selected genera (identified from the RF/XGB extreme-class models) in Bottom versus Top tiers, again grouped by FungalTraits guild. Bars show mean relative abundance of each genus within guilds and tiers, highlighting genera that increase or decrease between Top and Bottom trays.

To link these guild-level trends back to the specific taxa highlighted by the machine-learning models, we overlaid ML-selected genera onto the FungalTraits guild classifications and re-examined their relative abundances by performance tier (Fig. 6b). Within the ML-selected subset, Top trays were enriched in soil saprotrophs and mycoparasites and relatively depleted in ECM and plant-pathogenic genera. For example, Top trays contained substantially more ML-selected soil saprotrophs and mycoparasites (driven largely by *Apiotrichum* and *Hyphodiscus/Simplicillium*) and fewer ML-selected plant pathogens and dung saprotrophs. At the genus level, ECM *Thelephora* and the soil-saprotrophic yeast *Apiotrichum*, which has been proposed as a beneficial rhizosphere fungus (Kumla et al., 2020; Sousa et al., 2012), showed higher relative abundance in high-performing trays, whereas ECM *Rhizopogon*, the plant-pathogenic *Pseudoarthrographis* and dung saprotroph *Lasiobolidium* were concentrated in low-performing trays. These analyses provide a guild-level interpretation of the machine-learning results and identify fungal community patterns associated with planting-stock quality.

## 4. Discussion

This study shows that operational root-zone management in containerised pine production is associated with coordinated shifts in the root-associated mycobiome and planting-stock quality. The main contribution of this work is not simply that individual inputs affected seedlings directly, but that management combinations were reflected in both composite seedling performance and fungal community structure. Across the experiment, the root mycobiome retained a stable ectomycorrhizal backbone while showing treatment- and performance-associated shifts in ECM genera and lower-abundance saprotrophic, endophytic, mycoparasitic and pathogen-associated guilds. Together, these results support the idea that nursery-conditioned root mycobiomes can provide ecological information about planting-stock condition before planting.

Viewed through a plant–soil feedback framework, the containerised nursery plug can be interpreted as a managed root-zone ecosystem rather than a passive growth medium. This interpretation is consistent with plant–soil feedback theory, which emphasises that plant-associated biotic communities can feed back positively or negatively on plant growth (Bever *et al*., 1997; van der Putten *et al*., 2013), and with work showing that feedback concepts can bridge natural and agricultural systems (Ehrenfeld *et al*., 2005; Mariotte *et al*., 2018). More broadly, the shift from bare-root to containerised nursery production does not remove belowground biological complexity; rather, it concentrates microbial assembly within a confined and highly managed root-zone environment, consistent with evidence that nursery inputs can shape seedling growth and nutrient status, and fungal community structure (Carrillo et al., 2011; Rikala et al., 2004; Stuart *et al*., 2023). Because this experiment measured associations at nursery dispatch, rather than reciprocal feedback effects after reintroduction or outplanting, these patterns should be interpreted as feedback-compatible microbial conditioning rather than direct proof of plant–soil feedback causality.

Within this framework, the treatment comparisons are best interpreted as operational levers that modified the root-zone environment through different ecological pathways. Granular thiophanate-methyl/etridiazole showed the clearest association with higher seedling growth and root-system quality, consistent with activity against damping-off and root-disease complexes in forest nurseries (Cram, 2023), while the accompanying shifts in pathogen-associated and mycoparasitic guilds may be biologically relevant given the effects of damping-off and root-rot pathogens on seedling quality and survival, even though direct disease symptoms were not measured (Lamichhane et al., 2017; Reglinski et al., 2009; Won et al., 2018). The contrast between granular incorporation and liquid application suggests that timing, placement and persistence of crop-protection inputs within container media may matter, consistent with context-dependent responses to etridiazole/thiophanate-methyl mixtures and pesticide movement under container irrigation systems (Dumroese et al., 1990; Juntunen and Kitunen, 2003). Biochar, by contrast, shifted the root mycobiome without improving seedling growth, suggesting a substrate- or habitat-modification effect; biochar can provide habitable pore space for fungal hyphae (Jaafar et al., 2014), alter growing-media properties (Kaudal et al., 2016; Nemati et al., 2015), and produce context-dependent pine seedling responses under different fertilisation regimes (Dumroese et al., 2018; Eskandari et al., 2019). Finally, the maintenance of seedling performance under 75% fertigation indicates that moderate reductions in nutrient input may be possible without compromising planting-stock quality, aligning with studies showing that targeted fertilisation can maintain seedling morphology and nutrient status while improving nutrient-use efficiency and reducing leaching losses (Dumroese et al., 2005; Juntunen et al., 2002). Together, these responses support the broader interpretation that integrated root-zone management, rather than any single input alone, shapes the balance between seedling performance and fungal community structure. This combined treatment structure is central to the study’s contribution because operational nurseries do not apply substrate amendments, crop-protection regimes and fertigation in isolation, extending earlier work on fertiliser, fungicide, compost-tea amendment and biostimulant strategies in nursery production (Smaill and Walbert, 2013; Otero et al., 2020; Stuart *et al*., 2023; Chowdhury *et al*., 2026).

Fungal community profiles contained more predictive capacity towards the CPI class than the management treatments alone, suggesting that a management label records the treatment applied, whereas the mycobiome reflects the combined outcome of substrate properties, nutrient supply, seedling growth and microbial interactions within the root plug. This interpretation is consistent with broader evidence that plant-associated microbial communities are shaped by host state and environmental conditions (Trivedi et al., 2020), and with the use of microbiome profiles as biological indicators of ecosystem condition, although such indicators do not by themselves establish mechanism (Romero et al., 2025). The machine-learning models like those used here should therefore be interpreted as a diagnostic benchmark rather than evidence that individual fungal taxa caused higher CPI. The strongest ecological signal in the mycobiome data was the rebalancing of fungal guilds around a persistent ECM-dominated backbone. This pattern supports the premise that nursery-conditioned root-zone status is better described by relative shifts among functional guilds rather than by the abundance of any single genus. ECM fungi are central to conifer nutrient acquisition and can contribute to stress tolerance, but their effects depend on community composition rather than ECM presence alone (Colpaert et al., 1999; Parke et al., 1983; Smaill and Walbert, 2013). Root endophytes and dark septate fungi can also influence host performance and stress responses, although their effects are often context dependent (Mandyam and Jumpponen, 2005; Rodriguez et al., 2009). Saprotrophic fungi contribute to decomposition and nutrient cycling, while mycoparasitic fungi can suppress other fungi through antagonism or direct parasitism (Leifheit et al., 2024; Mukherjee et al., 2022). Because guild assignments are inferred rather than directly measured, these patterns should be interpreted as ecological indicators of root-zone status rather than confirmed functional mechanisms (Põlme et al., 2021).

This guild-level signal was expressed through turnover among major ECM genera as well as shifts in lower-abundance fungal groups. High- and low-performing CPI groups differed in the relative abundance of major ECM genera, suggesting turnover within the dominant symbiotic guild rather than a change in overall ECM dominance. This ECM turnover is ecologically relevant because *Rhizopogon rubescens* has been associated with foliar nutrient status and persistence after outplanting in radiata pine (Smaill and Walbert, 2013; Walbert et al., 2008), whereas *Thelephora terrestris* can be a strong root coloniser with a different competitive strategy in pine systems (Moeller and Peay, 2016). At the same time, lower-abundance fungal groups provide examples of how the guild-level signal may extend beyond ECM turnover, including changes in saprotrophic, endophytic and mycoparasitic groups linked to decomposition, host stress responses and fungal antagonism (Mašínová et al., 2017; Leifheit et al., 2024; Rodriguez et al., 2009; Mukherjee et al., 2022). *Apiotrichum* can be viewed as one candidate example within this broader pattern: it was enriched in Top-CPI trays and associated with higher biomass in a previous radiata pine nursery study (Chowdhury et al., 2026), a role supported by culture-based assays of soil yeasts, including *Apiotrichum scarabaeorum*, which have reported plant-growth-promoting traits such as IAA production (Kumla et al., 2020). These observations identify hypotheses for future testing but do not establish function in this experiment; guild assignments remain predicted ecological roles rather than direct functional measurements (Põlme et al., 2021).

Together, these findings position the containerised nursery plug as a managed ecological interface where production inputs, seedling performance and root-associated fungal communities intersect. The persistence and field relevance of these fungal community patterns remain unresolved. However, it is likely, based on published research, that they continue to have a strong effect. Nursery-associated ECM communities of radiata pine have been shown to persist for up to two years after outplanting, with *R. rubescens* among the most persistent taxa (Walbert et al., 2008). Similarly, reduced nursery fungicide exposure has been associated with improved radiata pine survival and diameter growth for six years after planting, even after treatment-related ECM differences were no longer detectable (Smaill et al., 2020). Integrating composite seedling traits with mycobiome profiling therefore provides a practical framework for evaluating biological dimensions of planting-stock quality before outplanting, while controlled inoculation, reintroduction and field experiments will be needed to determine whether the fungal community patterns observed here contribute to seedling performance or mainly indicate favourable root-zone conditions, following approaches that have used culture collections or defined microbial communities to test microbiome function in other plant systems (Bai et al., 2015; Castrillo et al., 2017).

## Statements and Declarations

### Ethics approval and consent to participate

Not applicable. This study did not involve human participants or animal experiments requiring ethical approval.

### Data availability

Raw ITS2 amplicon reads have been submitted to the NCBI Sequence Read Archive under BioProject accession PRJNA1535522.

### CRediT authorship contribution statement

**Jamil Chowdhury:** Methodology, Investigation, Data curation, Formal analysis, Visualization, Writing – original draft, Writing – review & editing. **Nathan Milne:** Resources, Investigation, Writing – review & editing. **Melanie Wade:** Resources, Investigation, Writing – review & editing. **Bronwyn Thuaux:** Investigation, Writing – review & editing. **Phil Green:** Supervision, Funding acquisition, Writing – review & editing. **Ian Last:** Supervision, Funding acquisition, Writing – review & editing. **John Senior:** Supervision, Funding acquisition, Writing – review & editing. **Angus J. Carnegie:** Supervision, Funding acquisition, Writing – review & editing. **Ian C. Anderson:** Supervision, Funding acquisition, Writing – review & editing. **Tarryn Turnbull:** Supervision, Funding acquisition, Writing – review & editing. **Krista L. Plett:** Supervision, Project administration, Funding acquisition, Formal analysis, Writing – review & editing. **Jonathan M. Plett:** Methodology, Investigation, Supervision, Project administration, Funding acquisition, Formal analysis, Writing – original draft, Writing – review & editing.

## Acknowledgement

We acknowledge the significant financial contributions from industry members and NIFPI’s commitment to advancing Australia’s forest and wood products industry through innovation in areas such as forest management, timber processing, and the bio-economy.

