## Supplementary material for "Root-zone modelling frameworks identify this managed ecological interface as a predictor of pine seedling quality": Fig S1 and S2

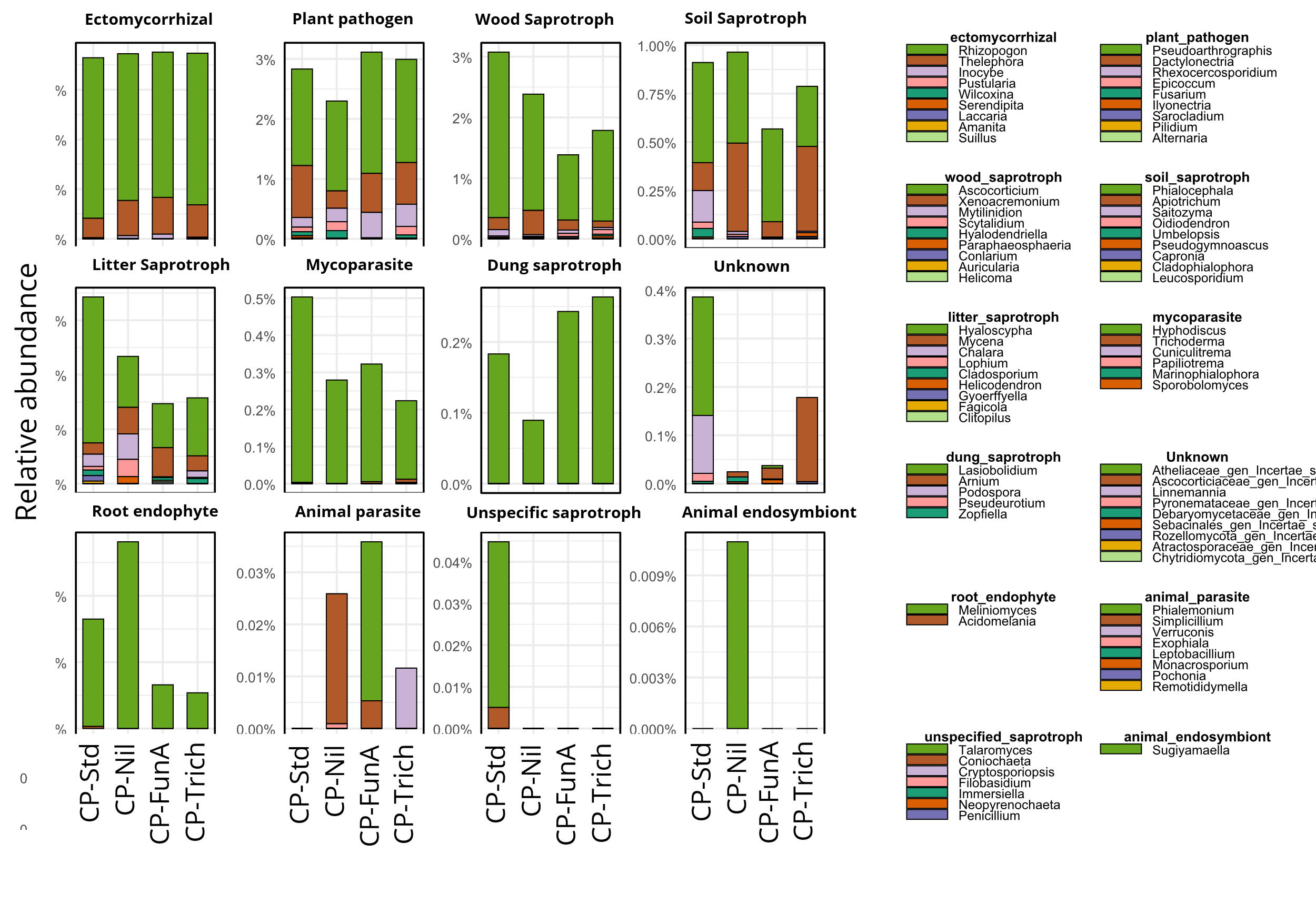

**Figure S1. Effect of crop-protection regime on functional guild composition of the root mycobiome.** Relative abundance of fungal functional guilds and dominant genera in 12-month seedlings exposed to alternative crop-protection regimes (standard programme, CP–Nil, CP–FunA drench, CP–Trichoderma drench), with substrate amendment, fertiliser and fertigation held constant. Panels and colours are as in Figure 3. Functional guilds were assigned using the FungalTraits database.

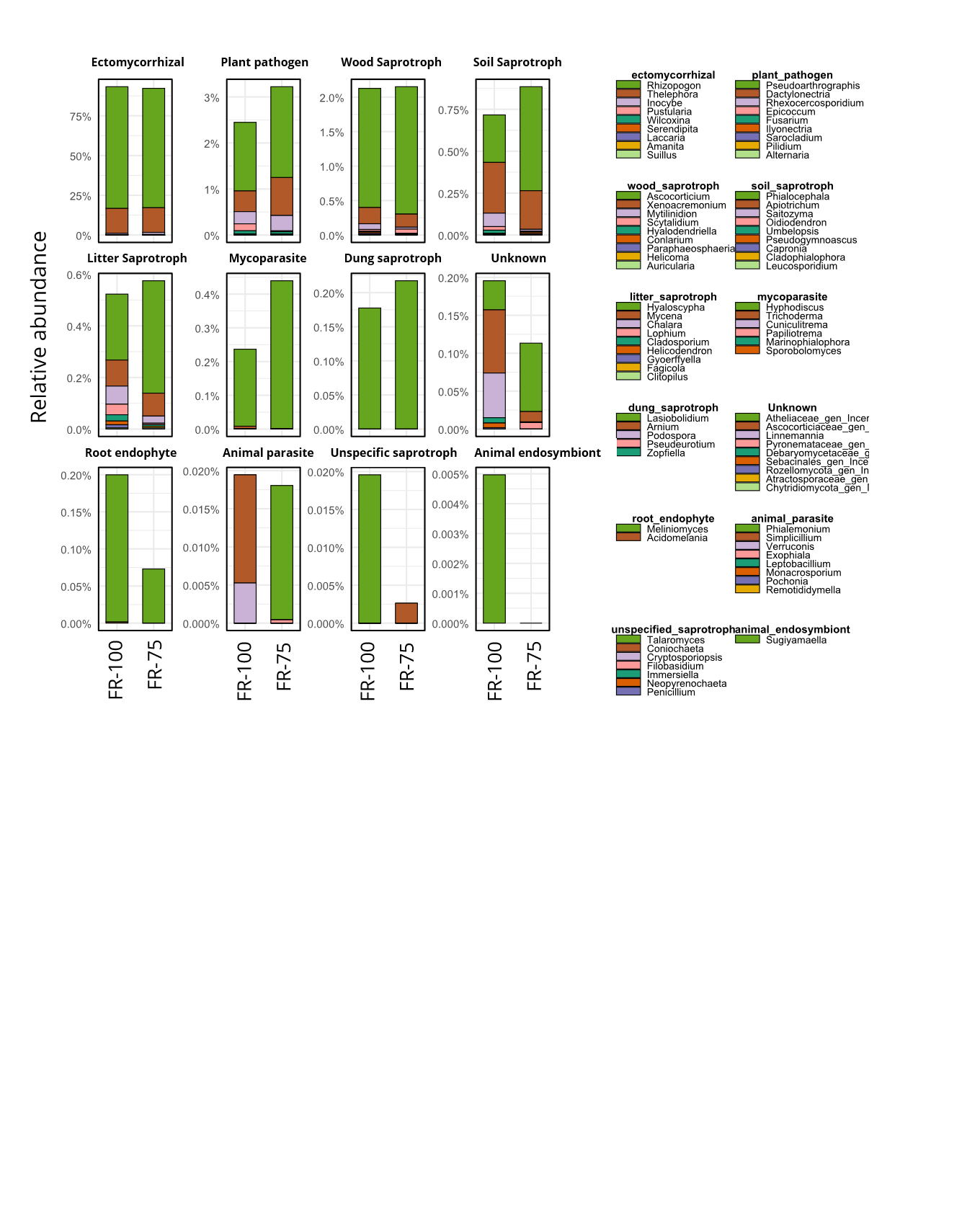

**Figure S2. Effect of fertigation rate on functional guild composition of the root mycobiome.** Relative abundance of fungal functional guilds and dominant genera in 12-month seedlings receiving 100% (FR-100) or 75% (FR-75) of the recommended fertigation rate, with substrate amendment and crop-protection regime fixed. Panels and colours are as in Figure 3. Functional guilds were assigned using the FungalTraits database.
